# Transcriptional regulation of structural and functional adaptations in a developing adulthood myocardium

**DOI:** 10.1101/2020.11.25.397885

**Authors:** Sini Sunny, Anil Kumar Challa, Joseph Barchue, Muralidharan T. Ramamurthy, David K Crossman, Steven Pogwizd, Senthilkumar Cinghu, Namakkal S. Rajasekaran

## Abstract

The development of the heart follows a synergic action of several signaling pathways during gestational, pre- & postnatal stages. The current study aimed to investigate whether the myocardium experiences transcriptional changes during the transition from post-natal to adult hood stages. Herein, we used C57/Bl6/J mice at 4 (28-days; post-natal/PN) and 20 weeks (adulthood/AH) of ages and employed the next generation RNAseq (NGS) to profile the transcriptome and echocardiography analysis to monitor the structural/functional changes in the heart. NGS-based RNA-seq revealed that 1215 genes were significantly upregulated and 2549 were down regulated in the AH versus PN hearts, indicating a significant transcriptional change during this transition. A synchronized cardiac transcriptional regulation through cell cycle, growth hormones, redox homeostasis and metabolic pathways was noticed in both PN and AH hearts. Echocardiography reveals significant structural and functional (i.e. systolic/diastolic) changes during the transition of PN to adult stage. Particularly, a progressive decline in ejection fraction (EF) and cardiac output was observed in AH hearts. These structural adaptations are in line with critical signaling pathways that drive the maturation of heart during AH. Overall, we have presented a comprehensive transcriptomic analysis along with structural-functional relationship during the myocardial development in adult mice.

## 1. Introduction

Beginning with the gestation, the heart undergoes a multitude of transcriptional and translational mechanisms to gain morphological and functional maturation (1). In a developing embryo, the heart is the first functional organ experiencing cell proliferation, vascular growth and development of an electrical conduction system (2). Programmed regulation of myocardial transcriptome at a given time (i.e. age) is crucial for the normal growth and function of the heart; Otherwise, development may culminate in serious cardiovascular outcomes (3). A close association between age and gene regulatory mechanisms facilitates constant signals for the developing heart. Maintaining a healthy growth and functional adaptation in the heart relies on various factors that directly or indirectly influence the structural and electrical conduction properties (4). Several studies have reported that conditions such as hypertension, atherosclerosis, diabetes, ischemia/reperfusion and altered circadian rhythm are coupled with pathological remodeling and dysfunction of the heart (5-7). Many of these events are also connected with altered transcriptional regulation and differential expression of several of the genes involve in cardiac structure and function. Nonetheless, actual transcriptional changes and gene regulatory networks that drives the development of post-natal to adulthood heart remain elusive.

Several “gene expression” and “omics” studies have reported the changes in cardiac development (8-10). Furthermore, most of these studies focused either on pre-natal development or on age-associated cardiac remodeling (11). Therefore, it is crucial to understand how transcriptional regulation is aligned with the structural and functional adaptation of the heart during the transition from post-natal to adulthood stages. Employing advanced technologies, such as next generation sequencing (NGS), to map out the transcriptome changes, and exploring the gene regulatory networks to align with the structural and functional adaptations of the post-natal heart would be informative.

Understanding the transcriptional mechanisms and gene regulatory networks that are aligned with structural and functional adaptations during adulthood are crucial for maintaining long-term cardiac health. Therefore, in the current study, we investigated the changes in the gene regulatory networks in post-natal and adult hearts, and explored the relationship between transcriptional profiles with the myocardial structure and function. We used NGS-based RNA sequencing, Ingenuity Pathway Analysis (IPA) and echocardiography to reveal the structural (size, volume & mass) and functional (systolic & diastolic) properties in a post-natal (28 days) and adulthood (4.5 months old) C57/Bl6/J mice. The results of this study established a novel connection between transcriptional changes and myocardial adaptation during adulthood.

## 2. Materials and Methods

### 2.1. Animals

Adult C57BL/6 J mice (wild type) at 4 (28 days; Post-natal/PN) and 20 weeks (Adulthood/AH) of age were used for analyzing the myocardial transcriptome and structural and functional remodeling (non-invasive echocardiography). Animals were maintained under hygienic environment with a 12 h day/night cycle, and provided standard animal chow and water *ad libitum*. All the animal studies including non-invasive cardiac imaging were conducted in accordance with the Guide for Care and Use of Laboratory Animals developed by the National Research Council at the National Institutes of Health (NIH). The Institutional Animal Care and Use Committee (IACUC#14-10160) at the University of Alabama at Birmingham has approved the study.

### 2.2. Next generation RNA sequencing

Total RNA was isolated from PN (4 weeks) and AH (20 weeks) old C57/Bl6/J mouse hearts (n= 4-6/group) using the RNeasy Mini Kit (Qiagen, Cat.74106) according to the manufacturer’s instructions. The purity of RNA was confirmed using Bio-analyzer and intact poly(A) transcripts were purified from total RNA using oligo(dT) magnetic beads and mRNA sequencing libraries were prepared with the TruSeq Stranded mRNA Library Preparation Kit (Illumina, RS-122-2101, RS122-2102). A D1000 Screen Tape assay (Agilent, Cat.5067-5582/3) was used with the 2200 Tape Station (Agilent Technologies) to quantify purified libraries. cBot was used to apply 18pM of the sequencing library to a TruSeq v3 flowcell (Illumina), and the TruSeq SR Cluster Kit (Illumina, Cat. GD-401-3001) was used for clonal amplification. Next, the flowcell was placed in the HiSeq. 2000 and underwent a 50 cycle single read sequence run with TruSeq SBS Kit v3-HS reagents (Illumina, Cat. FC-401-3002). The raw sequence reads were aligned to the reference genome using STAR (version 2.7.3a). Following alignment, transcript abundances were calculated using HTSeq-Count (version 0.11.3). The counts were then analyzed in DESeq2 (version 1.26.0), which normalized the signal and determined the differentially expressed genes (12).

### 2.3. Ingenuity Pathway Analysis (IPA)

Differentially expressed genes (DEGs) were analyzed using Ingenuity Pathway Analysis (IPA) software (Qiagen, Hilden, Germany) to determine over-represented canonical pathways and potential upstream or downstream regulators. The cut-off criteria used to carry out analyses were fold change (≥ ±2) & p-value (< 0.05). IPA generated the gene list for each disease annotations based on Z score (12). DEGs were then plotted for heat maps based on fold change with R studio package “pheatmap” (Version 1.2.5033). We also performed PANTHER analysis to examine the gene ontology enrichment and identify potential targets and unique interactions (*pantherdb.org*).

### 2.4. RNA isolation and real-time PCR validation

Total RNA was isolated from 10-20 mg heart tissue (RNeasy kit, Qiagen) of PN and AH mice (n≥6 in each group). After assessing the purity using a Nano Drop ONE^c^ (Thermo Scientific, Waltham, MA), 1.25 μg RNA was reverse transcribed to synthesize cDNA (QuantiTect Kit; Qiagen #205313) according to the manufacturer’s instructions. Using a QuantiTect SYBR Green PCR kit (Qiagen, Cat: 204145) and a 10 μl real-time qPCR reaction (with 30–37.5 ng cDNA template and 1.0 pmol of specific primer sets for targeted genes; Supplementary Table, S1) qPCR was performed using a Roche Light Cycler 480 (Roche Life Science). Copy numbers of cDNA targets were quantified using Ct values, and mRNA fold changes were calculated by normalization to the Ct of housekeeping genes *Arbp1* or *Gapdh* according to the 2^-ΔΔCt^ method (13).

### 2.5. Non-invasive echocardiography imaging of myocardium

Two-dimensional, Doppler and M-mode echocardiography acquisitions were performed in mice at respective ages [4W (n=18) & 20W (n=12)] with a high-resolution ultrasound system (Vevo2100, Fujifilm Visual Sonics Inc., Ontario, Canada) equipped with 38-MHz mechanical transducer (14). Animals were anesthetized (1.5–2.0% of isoflurane), placed on a warming platform (37°C) and the hair on the chest was shaved off to minimize signal interference. The heart rate, ECG signals and respiration were recorded, and the body temperature was monitored. Aqua sonic 100 gel was applied to the thorax area to optimize visibility of the cardiac chambers. Parasternal long-axis, parasternal short-axis and apical four-chamber, two-dimensional views were acquired. Cardiac structure, function (systolic & diastolic), and Tissue Doppler images were analyzed using Vevo lab 3.1 software.

### 2.6. Statistics

Fold changes in gene expression were calculated according to the 2^-ΔΔCt^ method and data are presented as mean ± SEM (n = 3-6/group). Basal comparisons between PN and AH mice were made using an unpaired t-test. All statistical analyses were performed using Graph Pad Prism-7 software with significance set at p<0.05.

## 3. Results and Discussion

### 3.1. Identification of differential transcriptome profiles in the myocardium of post-natal (PN) versus adulthood (AH) mice

The genetic underpinnings of transcriptional changes in growing postnatal heart are unclear. Herein, we employed the next generation RNA sequencing (NGS RNASeq) and trans-thoracic echocardiography to investigate the transcriptional changes and structural-functional changes, respectively, that occur during post-natal development in C57/Bl6/J (wild type) mice at 4 (post-natal; PN) and 20 (adulthood; AH) weeks of age (Fig 1A). Post-autopsy analysis showed a decrease in heart to body weight ratio in AH hearts (Fig 1B & C). While the heart size appeared to be increased in AH mice (0.3 vs 0.5 cm), a decrease in HW/BW ratio (4.7±0.21 vs 3.4±0.53) suggested physiological adaptation in AH mice. These programmed changes are crucial during the AH stage as this will determine the fate of cardiac health at later ages. Events like overnutrition or undernutrition during AH might profoundly influence cardiac pathophysiology with aging. Hierarchical clustering of the transcriptome data sets displayed distinct grouping of genes belonging to 4 and 20 weeks of ages in the heat map, suggesting robust changes between the PN and AH stages (Fig 1D). Analysis of differentially expressed genes (DEGs) at different fold levels revealed interesting results (Fig. 1E). There were 1230 (at ≥1.5 fold), 671 (at ≥2 fold), 350 (at ≥3 fold), 165 (at ≥5 fold), 80 (at ≥10 fold), 35 (at ≥20 fold), 13 (at ≥50 fold), and 5 (at ≥100 fold) genes downregulated significantly (p<0.05) in AH vs. PN hearts. On the other hand, there were 692 (at ≥1.5 fold), 277 (at ≥2 fold), 137 (at ≥3 fold), 70 (at ≥5 fold), 26 (at ≥10 fold), 8 (at ≥20 fold), 3 (at ≥50 fold), and 2 (at ≥100 fold) genes significantly (p<0.05) upregulated in AH compared to PN hearts. qPCR analysis (n=3/group) of selected genes (*Timp4, Gsta1, C1s1, Aldob1, Mmp9, Nppb, Mmp2, Acta2, Myh6, Timp3*) validated the NGS data (Fig. 1F).

**Figure 1:**
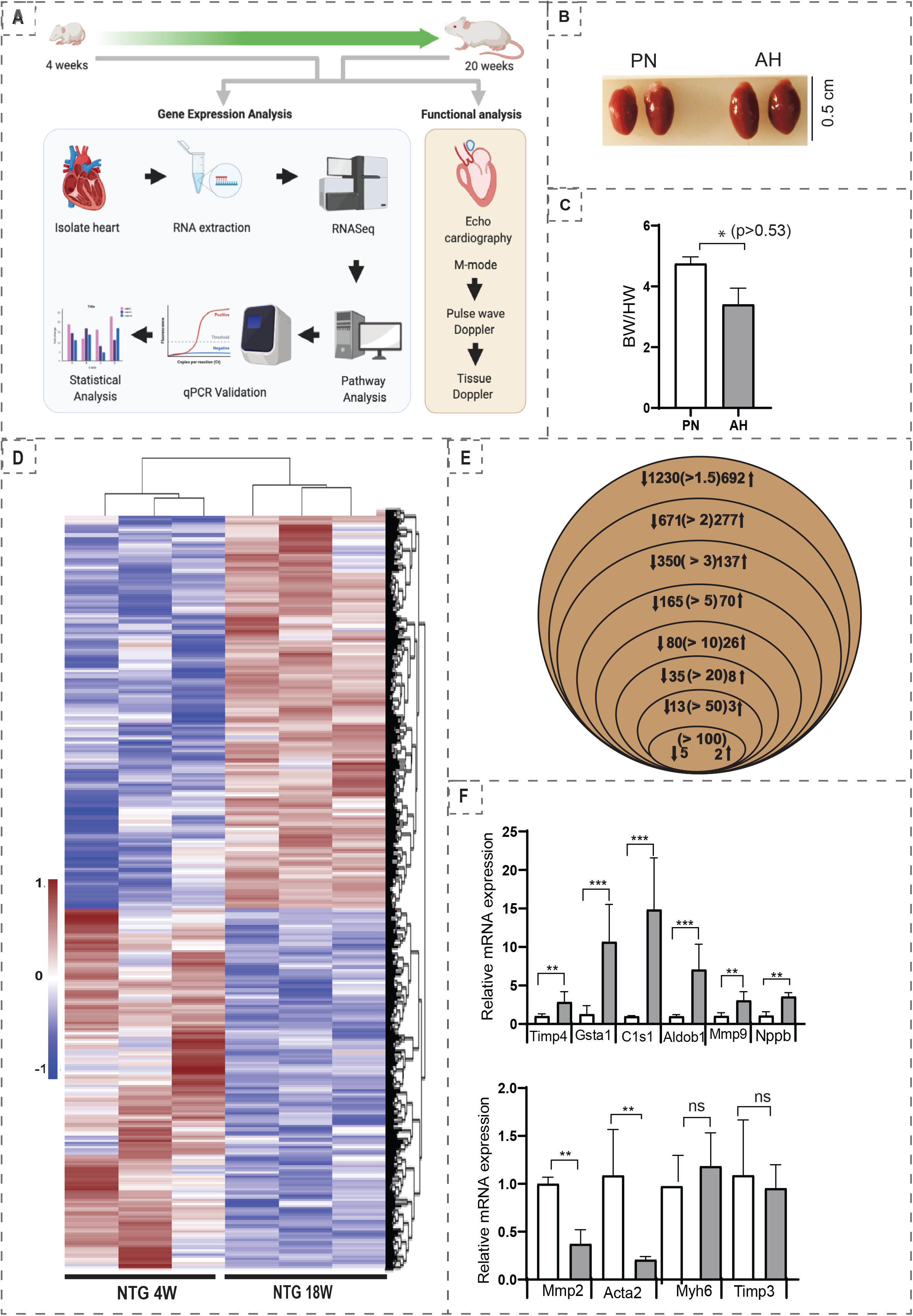
(A) Overall flowchart of the methodology. (B) Representative heart images of PN and AH mice (n=2/ group). (C) The heart-to-body weight ratios (HW/BW) measured at autopsy of PN and AH mice (n=4-5/group). (D) Venn diagram showing the number of significantly (P<0.05) upregulated and downregulated Differentially Expressed Genes (DEGs) based on fold change in PN vs. AH hearts. (E) Heat map of Z-score transformed row values for all significant (FC≥1.5, P<0.05) DEGs across the transcriptome. Each row represents a DEG wherein brown denotes upregulation; blue, downregulation; white, no change according to the color scale shown (side panel) (n=3 mice/group). (F) Real time qPCR validation of the most significantly changed cardiac remodeling genes (n=3 mice/group).

### 3.2. Altered transcriptional regulation revealed changes in the canonical pathways in AH hearts

Ingenuity Pathway Analysis (IPA), used to identify DEGs enriched in PN vs AH mouse hearts, showed enrichment of several altered biological functions that were statistically significant (p <0.05; Fig. 2). Upstream regulators identified from the list are mostly associated with cell cycle, embryonic development, cytoskeletal assembly, cardiomyocyte proliferation and cell-cell adhesion, cardiomyogenesis, NOS signaling, cellular assembly and several metabolic pathways, either directly or indirectly. Enriched transcriptomic pathways suggest continued developmental processes in the AH mouse hearts, in contrast to a “steady state” between 4 and 20 weeks. We also found that the levels of genes involved in various metabolic pathways like prostanoid and oleate biosynthesis were significantly increased in AH heart. Molecules involved in prostanoid pathway could modulate the progression of atherosclerosis and cardiac hypertrophy (15), which are also upregulated in AH hearts. These findings demonstrate physiological adaptations in AH hearts through measurable changes in the myocardial transcriptome.

**Figure 2:**
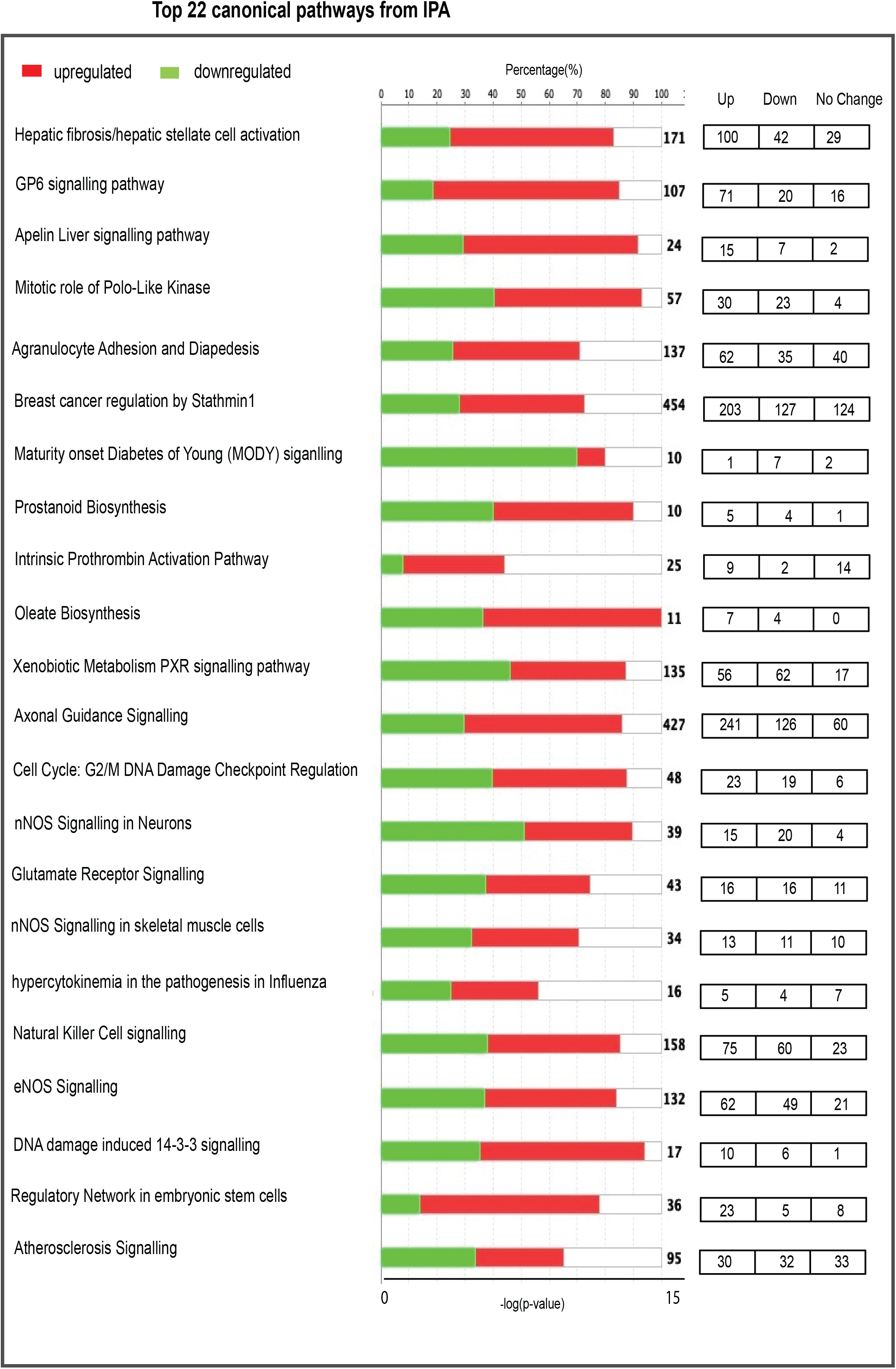
Top 22 canonical pathways significantly enriched in 4 week (post-natal; PN) vs. 20 week (adulthood; AH) mouse hearts. On the basis of the evidence stored in Ingenuity pathway analysis (IPA) library, the top enriched pathways are represented on the basis of z-score. Positive z-scores are represented in red (upregulated) and negative z-scores in green (downregulated). X-axis on top indicates percentage of genes present in the transcriptome to the total number of genes present in each selected pathway by IPA. The bottom X-axis represent −log p-value of each differentially expressed genes in the pathway. The transcriptome data were filtered to display genes with fold change greater than 2 at a significance of p<0.05.

### 3.3. Impaired cellular stress response, redox and nitric oxide signaling are evident in AH hearts

Redox signaling plays a vital role during the development and structural remodeling of growing myocardium (16). Here, we explored whether the transcriptional changes in AH heart influence myocardial redox signaling circuits. In this context, we were particularly interested in genes directly involved in redox homeostasis. We adopted hierarchical clustering of genes based on log fold change (FC>1.5) and extracted all the DEGs in response to cellular stress and vascular homeostasis (NO signaling) in the myocardium (Fig. 3). Overall, 1922 genes (Fig. 3A) were clustered with the log fold change +/-1.5 (FC>1.5, p<0.05). Among those, 1230 genes were downregulated and 692 genes were upregulated in the AH versus PN hearts. Most of the downregulated genes (Fig. 3B) observed in AH mice were involved in angiogenesis, growth hormone signaling, DNA replication, cytokine signaling, cell cycle, and apoptosis. Out of 692 upregulated genes, 15 were found to be specifically involved in nitric oxide (NO) signaling (Fig. 3C) and 50 were involved in Nrf2-mediated signaling (Fig. 3D) and xenobiotic metabolism (Fig. 3E). NF-E2-related factor 2 (Nrf2) is a key transcription factor, critical for cellular defense against oxidative and xenobiotic insults (17). Of note, upregulated xenobiotic metabolism, Nrf2 and NO signaling observed in AH hearts signifies the necessary redox and metabolic adaptations during adulthood. There was upregulation in AH hearts of some genes involved in oxidative deamination *(Maob*) and alcohol metabolism (*Aldh4a1, Aldh6a1, Aldh7a1*) as well as *Fmo5*, a phase 1 xenobiotic enzyme, suggesting the requirement of enhanced cytoprotective mechanism during this stage (18, 19). Upregulation of *Mgmt (1.7 fold)*, a gene crucial for genome stability was also observed, revealing an inherent defense mechanism by these hearts to prevent naturally-occurring mutagenic DNA mismatch, and errors during DNA replication and transcription (20). Genes involved in redox homeostasis by Nrf2 signaling were also upregulated in AH hearts. Key molecules involved in antioxidant response such as *Sod1 (1.2 fold), Sod2 (1.2 fold), Aox1 (2.4 fold), Cat (1.6)* were upregulated based on NGS fold change [Fig.5D]. Upregulation of key targets of *Nrf2, Nqo1 (2.3 fold), Gsta3 (1.9 fold), Gstm1 (1.4 fold), Gstm2 (1.8 fold), Gstm4 (1.6 fold)* shows adaptive response of the heart during postnatal development. Members of Dnaja family (i.e. *Dnaja1, Dnaja2, Dnaja3, Dnaj4*) and Dnajb family (i.e. *Dnajb1, Dnajb2, Dnajb3*) were also upregulated (Fig. 3D). Dnaj-dependent heat shock proteins are involved in the regulation of cell growth, apoptosis and protein homeostasis (21). In addition, some members in dnajb family are found to be decreased during obesity (22). Upregulation of genes responsible for NO signaling observed in AH hearts reveal changes in redox metabolism and energy requirements of the developing heart (23). However, NO production in turn depends on specific foods and diet, and it may vary during adverse conditions like undernutrition (24). Upon aging, anabolic pathways generating arginine decline, thereby impairing NO production (25). Likewise, a chronic practice of malnutrition or overnutrition in teenagers is likely to mimic aging conditions, which trigger pathologic stress in the heart. Therefore, our future studies will investigate the effects of premature aging in the heart of mice fed with high fat/sucrose diet. Overall, significant changes in the transcriptional regulatory networks related to sexual maturity, cardiac development, metabolism, and physiological adaptation take place in AH mice that facilitate structural and functional changes in the myocardium. The concerted action of these redox pathways reveals a properly coordinated endothelial physiology of the developing heart.

**Figure 3:**
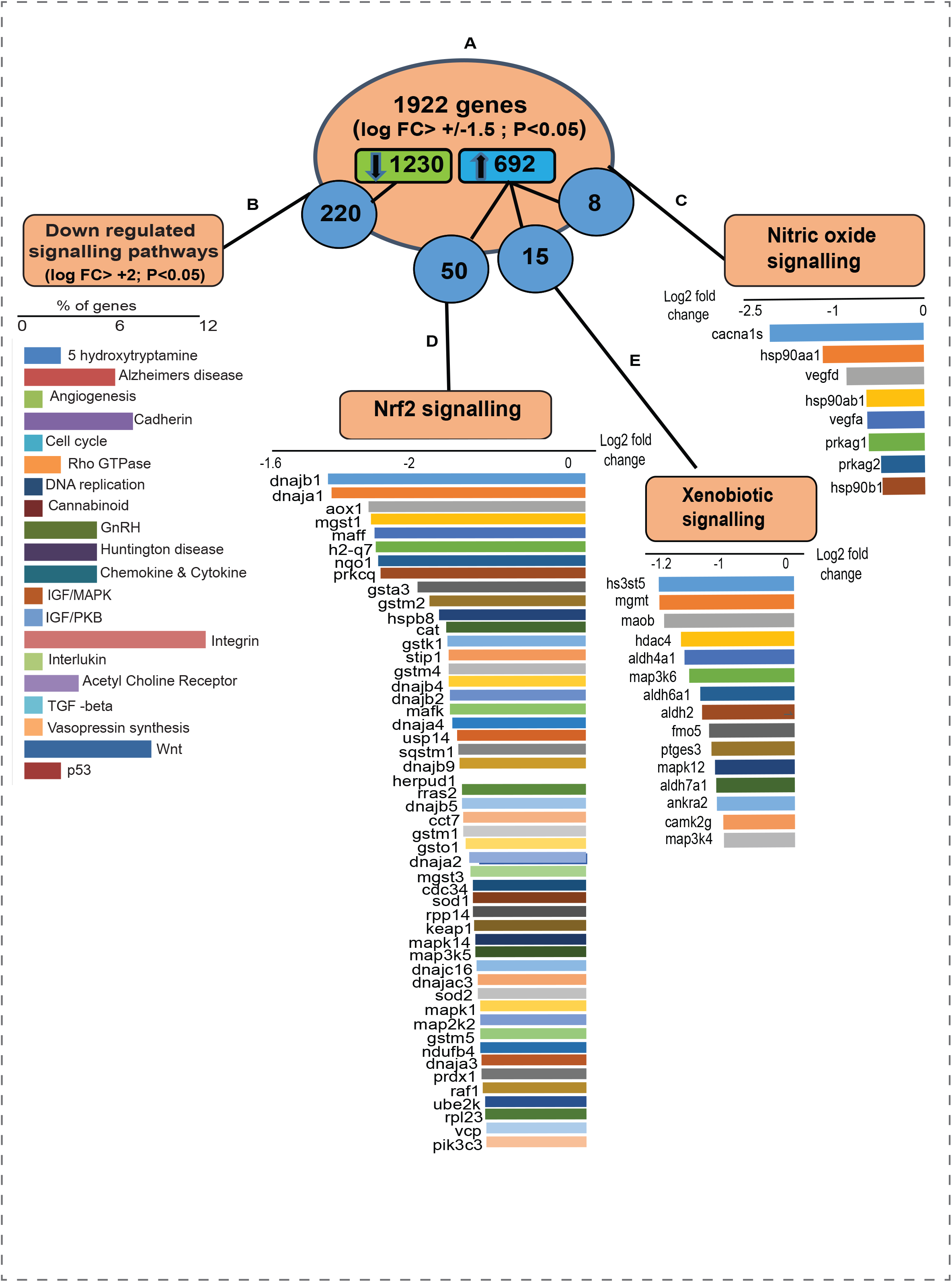
(A) 1922 genes were clustered based on log fold change +/- 1.5 with 1230 genes downregulated and 692 genes upregulated. (B) Downregulated genes (1230) were further clustered for fold change +/-2 and the enriched pathways were refined using PANTHER gene ontology analysis. Among 692 upregulated genes (P<0.05), 8 genes belong to (C) nitric oxide signaling, 50 belong to (D) Nrf2-mediated signaling and 15 belong to (E) xenobiotic metabolism pathway.

### 3.4. Most revealing transcriptome changes depict growth signaling in AH hearts

To elucidate the most revealing transcriptomic changes in AH hearts, hierarchical clustering of genes with a fold change of 10 and above (P<0.05) was employed to examine the up/down regulated genes in AH hearts (Fig. 4). Of note, Panther analysis of the upregulated genes displayed cadherin signaling as the primary event in AH hearts, indicating a critical developmental stage of multiple cardiac cell lineages and morphogenesis (26). The other major pathways represented by these upregulated genes included apoptosis, blood coagulation, β1/β2 adrenergic and cckr signaling and mitochondrial signaling pathways. On the other hand, the downregulated genes contributed to suppression of cadherin, integrin, Wnt, 5-hydroxy tryptamine degradation, TGF-beta, cannabinoid and Alzheimer’s disease signaling. These observations suggest that the enriched pathways constituted by both up- & down-regulated genes show the existence of co-coordinated signaling during late cardiac development. Taken together, significantly up- and -down regulated DEGs observed in AH mouse hearts demonstrate a dynamic transcriptional regulation during the transition from PN to adulthood.

**Figure 4:**
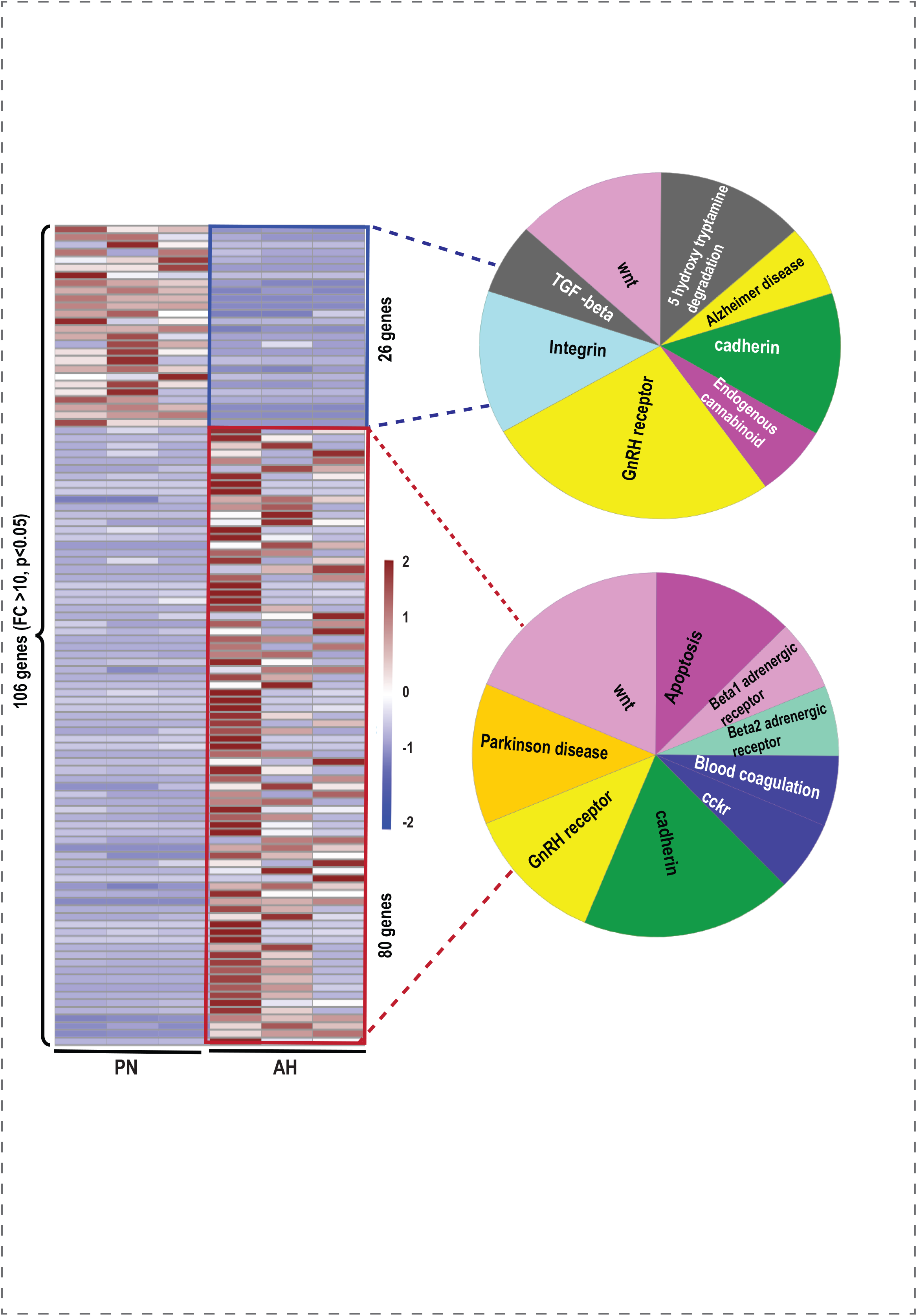
Heat map showing the distribution of 106 DEGs in PN vs. AH hearts (FC>10, P<0.05). Clustering annotations were omitted to plot the heat map in R studio. Pie diagram in the left and right side panel shows the upregulated (26 genes) and downregulated (80 genes) respectively. Enriched pathways of the genes in the two major clusters (up & down) were performed using PANTHER gene ontology analysis.

### 3.5. Identification of pathways related to cardiac development in the AH mice

Since hierarchical clustering analysis of genes (FC>10, Fig. 4) did not highlight key pathways involved in cardiac development and physiological adaptations, we employed IPA analysis to interpret gene ontology. Gene expression signatures distinguished by IPA have shown that not only the genes involved in the cardiomyocyte maturation phase but also the factors driving cardiac remodeling (physiological changes) have significant changes. Transcripts in one of the enriched pathways, sirtuin signaling (Fig. 5A) and its key molecules involved in mitochondrial bioenergetics as well as the cardiac developmental phase, were significantly altered. Upregulation of *Cpt1b and Mt-nd3* elucidates the role of highly coordinated myocardial triglyceride (27) and mitochondrial (28) metabolism. Decreased expression of *Dot1l* reflects the contribution of developmental arrest related to chamber dilation, cardiomyocyte survival and enhanced systolic function (29). Next, we observed the activation of Rho A signaling (Fig. 5B) that protects the developing heart from stress response (30). Decreased *Actg2* and *Igf1* reflect the developed cytoskeletal assembly and ventricular development in AH hearts (31, 32). Developing heart experiences an impaired *Cdc42* cell cycle signaling (Fig. 5C), through the downregulation of *Myl4, Myl7, and Myl 9* genes, which have been shown to play a key role in cardiogenesis (33). AH hearts also display decreased expression of *Fgfr2* (1.6 fold), *Fzd2* (1.5 fold) and *Gnai1* (1.1 fold) (Fig. 5D; p<0.05), which are involved in cardiac hypertrophy signaling (Refs). Hypertrophic growth of the heart is a “compensatory” response to hemodynamic stress (34). Downregulation of *Cdk1, Ccna2* and *Cdk6* (Fig. 5E) observed in AH heart indicate an arrest of cell cycle regulatory mechanisms and endothelial cell proliferation (35). Downregulation of *E2f2* observed in the transcriptome explains the decreased endothelial cell proliferation in AH hearts in the presence of ischemic insult (36). Downregulated paxillin signaling mediators (*Acta, Itgab, Itgb4, Pak3, Pik3cg*) (Fig. 5F) likely reflect fibronectin signaling that underlies the normal cardiac developmental features in AH hearts as reported earlier (37). Positive cell cycle regulators, such as cyclins, cyclin-dependent kinases (CDKs) are found to be upregulated in adult heart. In contrast, the present data show decreased gene expression of cell cycle mediators and the parallel upregulation of cardiac hypertrophy genes, suggesting the existence of an innate response by the cardiomyocytes to counteract the changes in the developing myocardium.

**Figure 5:**
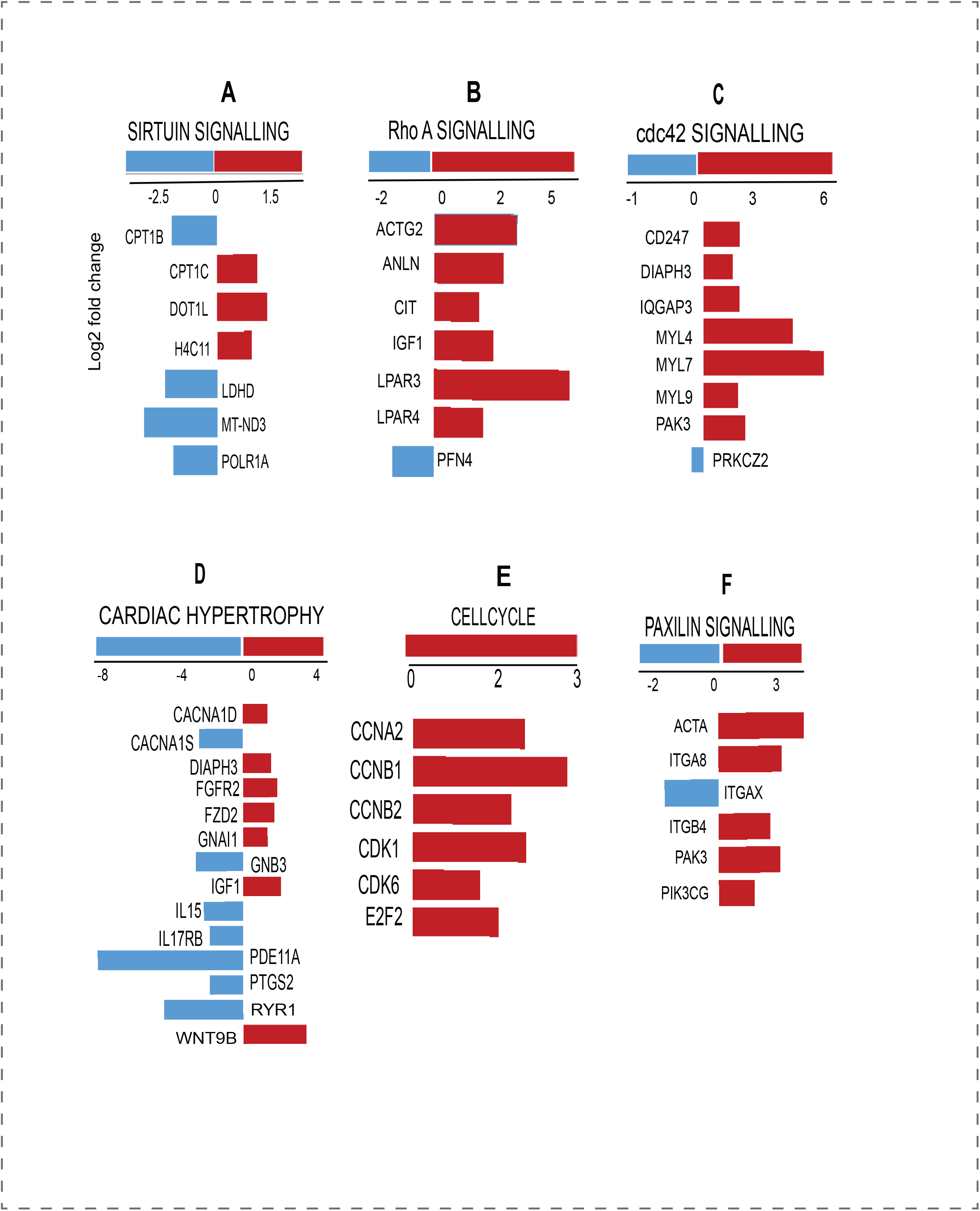
Enriched cardiac developmental pathways in 4 week (post-natal; PN) vs. week (adulthood; AH) mouse hearts. IPA analysis identified differential expression of several molecules (directly or indirectly) involved in cardiac developmental pathways in the transcriptome (FC>2; P<0.05; z-score >0.2). (A) sirtuin signaling, (B) Rho A signaling, (C) Cdc42 signaling, (D) cardiac hypertrophy signaling, (E) cell cycle and (F) paxillin signaling.

### 3.6. Structural and functional adaptations of adulthood myocardium

Using echocardiography (Vevo2100 Visual Sonics), we imaged and computed the structure and functional dynamics of the PN and AH hearts. M-mode analysis (Fig 6A) in parasternal short axis (PSAX) showed no significant changes in left ventricular volume (LVV), left ventricular interior diameter (LVID) and interventricular septum (IVS) of AH versus PN hearts. Significant changes in left ventricular mass (LV mass), ejection fraction (EF), and cardiac output (CO) were observed in the hearts of AH in relation to PN mice (Fig 6B). A negative correlation exists between LVV and LV mass among these hearts. A positive correlation between EF vs body weight (BW) and LV mass was noticed during the transition to AH stage (Fig 6C). Changes observed in the LV and CO directly rely on the age, which reveal that the development of myocardium continues through the AH stage. Assessment of LV diastolic function (Fig.6D & E), from mitral valve flow in apical 4-chamber views, revealed that there was a significantly decrease in mitral valve late (MVA) and early (MVE) filling rates. However, the overall LV diastolic function [MV (E/A) ratio] was comparable at both the stages. We noticed a positive correlation between the systolic (EF) and diastolic [MV (E/A) ratio] functions (Fig 6F). Tissue Doppler analysis at mitral valve (MV) (Fig. 6G-H) and tricuspid valve (TV) (Fig. 6I-J) levels exhibited no significant changes in velocity levels, which is an index of good cardiac health (38). Through *in vivo* assessment of the myocardium, using LV mass and cardiac output & EF, we noticed that there was a coordinated adaptation between structural and functional changes during the transition of PN to AH. During physiological growth, cardiomyocytes, endocardial cells, and epicardial cells coordinately respond to a variety of environmental and nutritional signals, which will have an impact on cardiac pressure loading, as well as electrical and hemodynamic forces. Transcriptomic changes observed in AH hearts reflect on its progressive structural and functional adaptations during that transition. Thus, maintaining appropriate nutritional practices and a stress-free environment during the transition to adulthood stage is crucial for stable growth and health of the myocardium.

**Figure 6.**
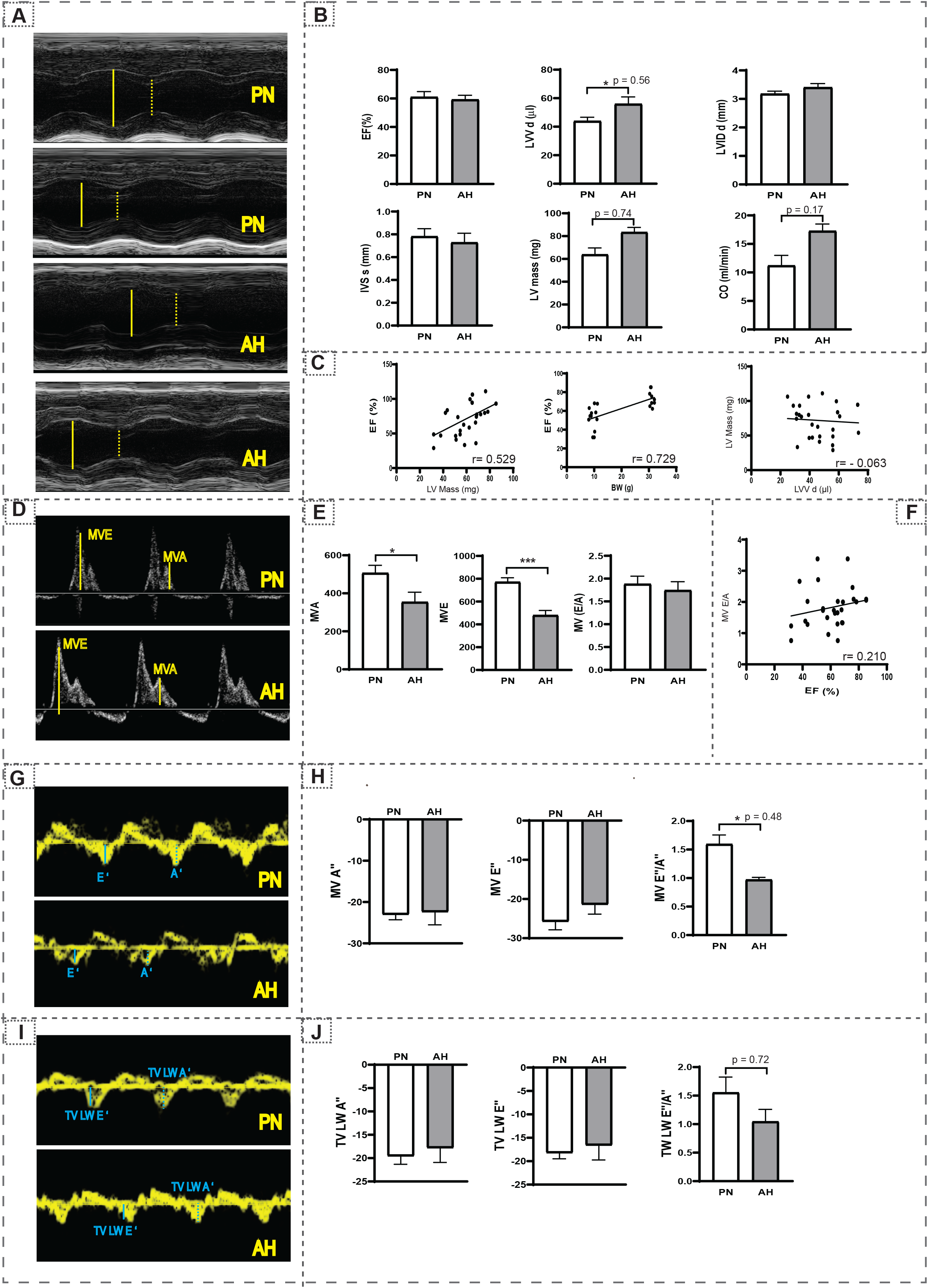
Echocardiography analysis in PN and AH hearts. (A) Parasternal short-axis (M-mode) echocardiographic images obtained for PN and AH mice (n = 12-18/group) using a high-resolution (38 MHz) ultrasound. The LV cavity was labeled in diastole (solid yellow line) and systole (dashed yellow line). (B) Cardiac structural and systolic functional changes measured using longitudinal strain and m-Mode analysis in PN and AH mice. (C) Correlation analysis for LV mass with body weight and ejection fraction, LV mass with LVV. (D) Blood in-flow through mitral valve captured using Pulse wave Doppler in PN and AH mice. (E) The mitral valve movement (mm) during early atrial filling (MV E, solid yellow line) and late atrial filling (MV A, dashed yellow line) were used to measure the diastolic function (MV E/A ratio). (F) Correlation analysis for ejection fraction (EF) with diastolic function (MV E/A ratio). (G) Tissue Doppler images taken at apical 4-chamber views at the mitral valve level. (H) Peak velocities at early (E’, solid blue line) and atrial (A’, dashed blue line) were measured. (I) Tissue Doppler images taken at apical 4-chamber views at the tricuspid valve level. (J) Peak velocities at early (TV LW E’, solid blue line) and atrial (TV LW A’, dashed blue line) were measured.

In addition to genetic and biochemical factors, diet and life-style significantly alters gene regulation, thereby influencing the development of heart. Developmental stages of mammals include multiple stages - pre-natal, infancy, early childhood, late childhood, adolescence, early adulthood and old age. During each of these stages, cells and organs adapt to retain structural and functional properties through continued active transcriptional and translational processes. Our data demonstrate that distinct changes occur in the transcriptome and structure/function of the myocardium during the transition to adult stage. Overall results of the study and its implication in an enhanced understanding of cellular processes that are regulated during the transition from PN to AH.

## 4. Translational statement/impact

Insufficient or surplus nutrition and irregular eating habits result in impaired development and function of skeletal and myocardial muscles. While inadequate nutrition results in decreased myocardial mass and cardiac performance, overnutrition might induce chronic metabolic stress, which then leads to pathological cardiac remodeling. Our future directions will focus on understanding the impact of mal- and surplus-nutrition on myocardial transcriptome/metabolism and cardiac health during adulthood.

## Acknowledgement

Authors thank Mr. Arun Jyothidasan and Dr. Shanmugam Gobinath for assisting with the methodology for bioinformatics.

## Ethics Statement

This study was carried out in accordance with the recommendations of the Guide for the Care and Use of Laboratory Animals of the National Institutes of Health. The protocol was approved by the Institutional Animal Care and Use Committee (IACUC) at the University of Alabama at Birmingham.

## Funding Information

This study was peripherally supported by funding from NHLBI (2HL118067 and HL118067) and NIA (AG042860) and the start-up funds (for N.S.R.) by the Department of Pathology and School of Medicine, the University of Alabama at Birmingham, AL, and UABAMC21 grant by the University of Alabama at Birmingham, AL. No direct funding is available for this work.

## Author Contributions

The study was designed by NSR. Experiments were performed by SS and NSR. NSR and SS interpreted the data and wrote the manuscript. JB, DKC and CS carried out IPA analysis. AKC, DKC, MTR, SP and CS were involved in interpreting the data and critical discussions. All authors read and approved the final version of this manuscript.

## Data availability

All data generated for this study will be provided as open-access on NCBI Gene Expression Omnibus (GEO) (mRNA) with all coding scripts and quality assessment at will be uploaded and disclosed to the public access.

## Competing Interests

The authors have no competing interests to declare.

## Consent for publication

All authors verified the content and approved the final version for submission and publication

## Notes

### Competing Interest Statement

The authors have declared no competing interest.

