## Supplemental Table 1 for "Transcriptional regulation of structural and functional adaptations in a developing adulthood myocardium"

This file includes:

Table. S1. Complete list of primer sequences used for qPCR analysis

**Supplementary Table S1**

**Complete list of primer sequences used for qPCR analysis**

| Gene Symbol | Forward | Reverse |
|---|---|---|
| *Gsta1* | AGCCCGTGCTTCACTACTTC | TCTTCAAACTCCACCCCTGC |
| *Acta2* | CCCTGGAGAAGAGCTACGAAC | TTTCGTGGATGCCCGCTG |
| *Timp3* | GAAGCCTCTGAAAGTCTTTGTGG | ACATCTTGCCTTCATACACGC |
| *Timp4* | TGTGGCTGCCAAATCACCA | TCATGCAGACATAGTGCTGGG |
| *c1s1* | GGCCCACTTGTTCCCATCAG | TTGGTAGTGAGGGACCACCC |
| *Myh6* | GAGTGGGAGTTTATCGACTTCG | CCTTGACATTGCGAGGCTTC |
| *Arbp1* | GAGATTCGGGATATGCTGTTGG | CGGGTCCTAGACCAGTGTTCT |
| *nppb* | GAGGTCACTCCTATCCTCTGG | GCCATTTCCTCCGACTTTTCTC |
| *aldob1* | GAAACCGCCTGCAAAGGATAA | GAGGGTCTCGTGGAAAAGGAT |
| *mmp9* | GCTGACTACGATAAGGACGGCA | TAGTGGTGCAGGCAGAGTAGGA |
| *mmp2* | AGCGAGTGGATGCCGCCTTTAA | CATTCCAGGCATCTGCGATGAG |
| *Gapdh* | TGACCTCAACTACATGGTCTACA | CTTCCCATTCTCGGCCTTG |
